# Genetic Expectations in Inheritance: A Probabilistic Algebraic Framework

**DOI:** 10.1101/2025.06.12.659255

**Authors:** Padavinangady Nakul Bhat, Debnath Pal

**Affiliations:** Department of Computer Science and Engineering, Manipal Institute of Technology, Manipal Academy of Higher Education, Manipal, 576104, Karnataka, India; Department of Computational and Data Sciences, Indian Institute of Science, Bengaluru, 560012, Karnataka, India

**Keywords:** genetic expectations, probabilistic genotype distributions, algebraic genetics, inheritance simulation, multi-locus simulation, computational genetics

## Abstract

This study introduces a formal framework for modeling inheritance patterns based on an algebraic representation of genotype distributions. The approach defines genotypes as probability measures rather than discrete states, enabling the computation of *genetic expectations*, a probabilistic analogue that captures the expected distribution of offspring genotypes under Mendelian inheritance rules. This formulation permits constant-time simulation of multi-locus inheritance processes and provides a means of incorporating uncertainty and partial information in genotype data. A series of eleven illustrative examples is presented, encompassing both Mendelian and selected non-Mendelian mechanisms, including polygenic inheritance, uniparental disomy, and haplodiploidy. While the current framework does not model recombination, linkage, or phenotypic traits, it is designed to accommodate extensions to some complex chromosomal structures, such as those encountered in polysomic inheritance. The model reproduces classical inheritance outcomes and is intended as a computational and theoretical tool for genetic analysis in both research and pedagogical settings. Future development will focus on increasing biological scope and integrating empirical data sources.

**Highlights:**

- Genotypes modeled as probability distributions over allelic states.
- Genetic expectations summarize an individual’s contribution to off-spring.
- Expectations are mating-independent, enabling modular inheritance logic.
- Framework supports constant-time, multi-locus inheritance simulation.

## 1. Introduction

The concept of genetic algebras was introduced by Etherington [4] and further developed by Schafer [10], Holgate [7], and Gonshor [6] as a formalism for representing inheritance processes algebraically. These structures provide a mathematical basis for describing the transmission of genetic traits across generations and have been applied to both Mendelian and non-Mendelian systems. However, much of the early work in this area emphasized abstract algebraic properties, with limited focus on computational considerations or applicability to large-scale genetic data.

This study develops a simplified algebraic model of Mendelian inheritance that is oriented toward computational tractability and probabilistic representation. The model introduces the notion of *genetic expectations*, wherein genotypes are treated as probability distributions over allelic configurations. This formulation allows for the simulation of multi-locus inheritance processes with constant-time complexity and can accommodate incomplete or uncertain genotype information. These features are of relevance in the context of high-throughput genetic data and population-scale studies.

The proposed framework differs from classical gametic and zygotic algebras in that it maintains genotype-level representations across generational transitions, avoiding intermediate transformations [9]. The model supports a range of Mendelian inheritance patterns and is structurally compatible with extensions to more complex chromosomal systems, such as polysomic inheritance and polyploidy, under the assumption of independent assortment.

Certain biological simplifications are retained for tractability, including the absence of mating structure (implicitly assuming random mating), no recombination, and the exclusion of evolutionary forces such as selection or mutation. As such, the framework does not aim to model full biological realism, but rather to provide a tractable basis for simulation, data pre-processing, and instructional applications. Potential extensions include the incorporation of recombination, linkage, and phenotype prediction.

To evaluate the model’s utility, a series of eleven examples is presented, covering both classical Mendelian cases and a range of non-Mendelian systems, including sex linkage, chromosomal abnormalities, and polygenic inheritance. These serve to illustrate both the generality of the formalism and its practical limitations. The objective of this work is to contribute a computational model of inheritance that is grounded in algebraic principles while remaining suitable for integration into contemporary genetic analyses. The framework is intended to support both theoretical exploration and practical applications in genetic modeling.

## 2. Background and Related Work

This section provides an overview of foundational concepts and prior research relevant to our work. The structure and much of the content follow the treatment presented by Reed [9], which serves as the primary source for this background. While some original papers were also consulted to enrich the discussion, the majority of the definitions and key ideas summarized here are drawn from Reed’s comprehensive analysis. We would also like to mention [12], which, while tangentially related, directly inspired the notation for our model.

### 2.1. Baric Algebras

Genetic algebras were first formalized by Etherington [4]. By his definition, an *algebra with genetic realization* is any algebra *A* with an *n*-dimensional basis {*a*_1_, *a*_2_, …, *a*_*n*_} over an appropriate field, usually ℝ. Multiplication is given by

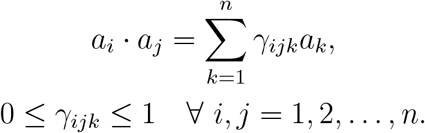

Additionally, if the algebra is *symmetric* [14], then

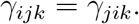

An element *x* ∈ *A* can be written as

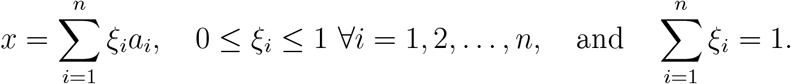

A *baric algebra* is a genetic algebra equipped with a non-trivial one-dimensional algebra homomorphism (weight function) *ω* : *A* →*k* to a field *k* [4].

Baric algebras are commutative, distributive, and generally non-associative [5]. The multiplicative powers of an element *x ∈ A* are defined as follows:

- **Principal powers:**

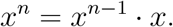
- **Plenary powers:**

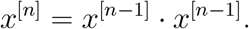

An *idempotent* element satisfies:

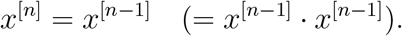

### 2.2. Interpretation of Baric Algebras

Baric algebras form the algebraic foundation for modelling genetic inheritance and adhere to Mendel’s laws.

Genetically, the *principal powers* of an element symbolize repeated mating between previous generation and the original generation, while the *plenary powers* represent self-mating within the previous generation.

An idempotent in this context represents a state of *genetic equilibrium*, where further generational iterations do not change the genetic composition [9].

The homomorphism *ω* : *A*→ *k*, known as the *weight function*, can be interpreted as a structure-preserving map that projects genetic traits into a scalar measure.

By carefully choosing *ω*, we can quantify certain desirable characteristics in an element. For example, let *A* = span {*D, d*}, where *D* is a dominant allele and *d* a recessive allele. Define:

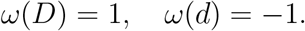

Then, for an element *x ∈ A*,

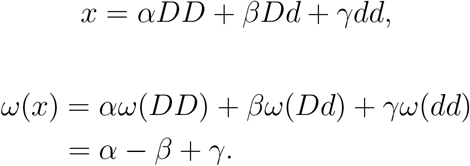

This expression quantifies the homozygosity of an element.

### 2.3. Train Algebras

In any algebra, for all elements, we can define a normalized polynomial called the *minimal polynomial m*_*x*_(*x*) of minimal degree such that it annihilates a single element in *A* [14]; i.e.,

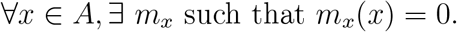

In baric algebras, we can extend the idea to a polynomial in principal powers, of minimal degree, that annihilates all elements of *A*. This polynomial is called the *rank polynomial* of *A* and is written as:

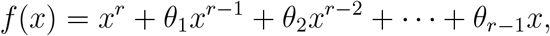

where *θ*_*p*_ is a homogeneous polynomial of degree *p* in the coordinates *ξ*_*i*_ of the generic element *x* ∈ *A*. The minimal degree *r* is called the *rank* of *A*. Additionally, if *θ*_*p*_ depends only on *ω*(*x*), then the baric algebra is called a *train algebra* [4].

### 2.4. Interpretation of Train Algebras

From the rank polynomial, we can derive the *rank equation* of *A*,

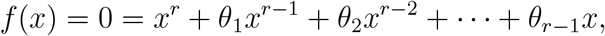

which gives:

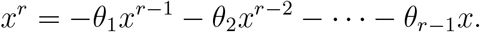

The rank equation provides a recurrence relation in principal powers, illustrating how the current genetic pool relates to previous genetic pools. As shown in [9], train algebras have uniquely determined weight functions.

### 2.5. Special Train Algebras and Genetic Algebras

In a train algebra *A* with a weight function *ω*, we define a special subalgebra

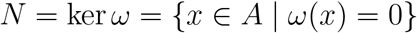

to be the kernel space of the weight function. We also define the powers of *N* as:

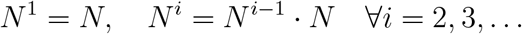

Etherington [4] defines a *special train algebra* as a train algebra where *N* is nilpotent and all powers of *N* are ideals of *A*. An ideal is a subspace that satisfies the property:

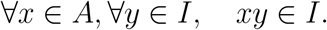

As Etherington observed, this definition classifies all gametic algebras as special train algebras. However, it overlooks some zygotic algebras, which also arise from these structures but are not accounted for.

To address this, Schafer [10] proposed a new class of algebras, relaxing the definition of special train algebras. A *genetic algebra* is a commutative baric algebra *A* such that the characteristic function |*λI −T*| of any transformation *T* = *αI* + *f* (*R*_*x*_, …, *R*_*x*_) ∈ *T*(*A*) depends only on *ω*(*x*_1_), …, *ω*(*x*_*n*_).

However, proving membership of an algebra according to Schafer’s definition is computationally intensive. Addressing this, Gonshor [6] provided an equivalent but more tractable formulation: a commutative finite-dimensional algebra *A* is genetic if there exists a basis {*a*_0_, *a*_1_, …, *a*_*n*_} such that

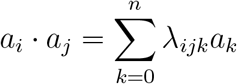

with multiplication constants satisfying:

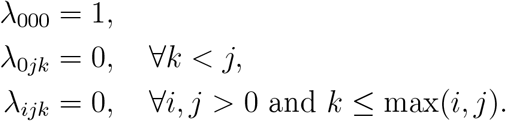

In his proof showing equivalence with Schafer’s definition, Gonshor showed that:

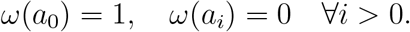

### 2.6. Interpretation of Special Train Algebras and Genetic Algebras

In Etherington’s framework, the kernel *N* = ker *ω* represents types that, while still capable of mating, are locked into a degenerative evolutionary path—effectively *non-recoverable* rather than strictly *sterile*. The nilpotency of *N* implies that repeated reproduction among these types leads to the eventual loss of viability, modelling a form of mutational decay. If all powers of *N* are ideals, then even mating with wild-type (viable) individuals fails to restore them: the offspring remain confined within *N*. This captures the idea of irreversible mutational drift—once a lineage enters *N*, it cannot revert to the wild-type.

Schafer’s definition, based on transformation algebras, implies that a system’s genetic evolution depends only on the *distribution of traits*, rather than on their specific identities. Holgate [7] proved that this condition holds under certain symmetry assumptions—symmetries that are prevalent in traditional Mendelian inheritance.

In Gonshor’s framework The basis element *a*_0_ represents the *wild-type* genotype, capable of viable reproduction, while the elements *a*_*i*_ with *i >* 0 represent mutants. The multiplication table implies:

- *a*_0_ *· a*_0_ = *a*_0_: wild types reproduce as wild types.
- *a*_0_ *· a*_*i*_ = *a*_*i*_: mating with a mutant preserves the mutation.
- *a*_*i*_ *· a*_*j*_ = *a*_*k*_: mutants produce more mutants.
- Repeated mating among mutants eventually leads to sterility.

This models irreversible degenerative mutation dynamics—analogous to *Muller’s ratchet*^2^—in which deleterious mutations accumulate in asexual populations, leading to reduced fitness and eventual sterility.

### 2.7. Bernstein and k-th Order Bernstein Algebras

Etherington’s Baric and Train algebras mainly study the properties of principal powers, which make more mathematic sense. However, genetically, an algebra based on the plenary powers makes more sense. Parallel to Etherington’s developments, S. N. Bernstein [3] studied some quadratic evolutionary operators that modelled inheritance using a genetic frequency simplex

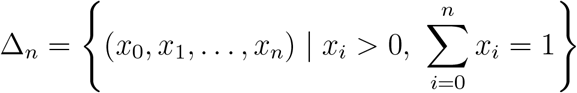

The operator Ψ maps such simplexes into themselves, symbolizing the next generation.

Bernstein focused on operators Ψ that satisfy the *stationarity condition* Ψ^2^ = Ψ, indicating that the population reaches equilibrium after one generation.

Later, Holgate [8] developed on Bernstein’s work and defined multiplication on a real vector space *V* with basis {*a*_0_, *a*_1_, …, *a*_*n*_} as

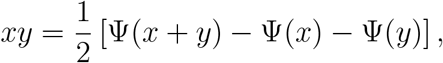

making *V* a commutative, non-associative algebra. A linear form *ω* : *V* →ℝ, defined by *ω*(*a*_*i*_) = 1 for all *i*, gives this a *baric* structure.

This framework yields the defining identity:

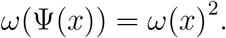

Based on this, we define a *Bernstein algebra* as follows. Let *A* be a finite-dimensional, commutative baric algebra over a field *k* with char(*k*) ≠ 2, and let *ω* : *A → k* be the weight function. Then *A* is called a *Bernstein algebra* if, for all *x ∈ A*, the plenary powers satisfy

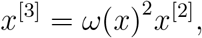

where *x*^[*n*]^ denotes the *n*-th plenary power of *x*.

Since many systems do not achieve stationarity after a single generation, Abraham [1] introduced a generalized version of Bernstein algebras, which considers equilibrium reached after *k* generations. This leads to the following definition:

Let *A* be a finite-dimensional, commutative baric algebra over a field *K* with char(*K*) ≠ 2, and let *ω* : *A*→ *K* be the weight function. Then *A* is called a *k-th order Bernstein algebra* if, for all *x* ∈ *A*, the plenary powers satisfy:

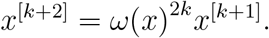

### 2.8. Interpretation of Bernstein and k-th Order Bernstein Algebras

Bernstein algebras abstract the dynamics of sexual reproduction, where genotype frequencies evolve quadratically across generations. The defining identity

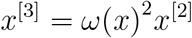

reflects an idealized scenario in which genetic equilibrium is reached after a single generation of mating. Here, *ω*(*x*) represents the reproductive success of genotype *x*, and plenary powers simulate repeated self-mating.

In contrast, many biological systems exhibit delayed stabilization due to factors such as recombination, mutation, or gene interactions. To capture these effects, *k*-th order Bernstein algebras relax the immediate equilibrium condition, requiring

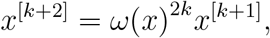

which implies that equilibrium emerges only after *k* generations.

### 2.9. Evolution Algebras

The algebras we discussed up to now mainly focused on Mendelian inheritance, and only accommodated some divergence. A recent development in the field, *Evolution algebras* were introduced by Tian [13]. They are specially designed to accommodate non-Mendelian inheritance such as asexual reproduction, uni-parental inheritance and other non-Mendelian systems.

An *evolution algebra E* over a field *K* is a commutative, (generally) non-associative algebra with a basis {*e*_1_, *e*_2_, …, *e*_*n*_} (called a *natural basis*) such that the product of any two distinct basis elements is zero:

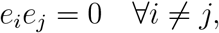

and the square of each basis element is a linear combination of basis elements:

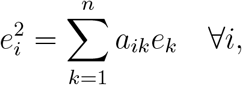

where *a*_*ik*_ ∈ *K* are called the *structure constants* of the evolution algebra.

### 2.10. Interpretation of Evolution Algebras

Evolution algebras offer a natural framework for modelling non-Mendelian inheritance patterns, particularly in systems where reproduction is asexual. Rather than relying on classical pairwise genetic interactions, these algebras describe inheritance by specifying how an individual’s type influences the types of its offspring.

In this setting, each basis element *e*_*i*_ represents a distinct genetic type (such as a genotype or an organism). The multiplication rule

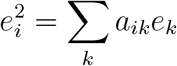

encodes the reproduction process: the coefficient *a*_*ik*_ represents the frequency or probability with which type *e*_*i*_ produces offspring of type *e*_*k*_. The requirement that *e*_*i*_*e*_*j*_ = 0 for *i* ≠ *j* reflects the absence of interaction between different types during reproduction, allowing only the modelling of asexual reproduction.

## 3. Model

Classical algebraic models of inheritance—including baric, train, Bernstein, and evolution algebras—offer a probabilistic foundation for describing genetic transmission, yet are primarily intended for abstract mathematical investigation. While these models effectively encode key features of Mendelian and some non-Mendelian inheritance patterns, their generality and structural complexity can limit their practical utility in computational settings.

The model presented here is explicitly constructed to support computational analysis of genetic inheritance. It provides a well-defined and interpretable framework for representing genotypic structure, evaluating inheritance dynamics, and reasoning probabilistically about allele distributions. By deliberately constraining the generality of the algebraic formulation, the model achieves several computational advantages, including ease of implementation, modularity in reproductive computations, and clarity in biological interpretation.

A defining component of the model is the concept of an individual’s *genetic expectation*, which represents a probabilistic summary of their genotype and quantifies their expected contribution to offspring. Importantly, this value can be computed independently of any mating partner, enabling efficient simulation of genetic crosses at scale. This independence also facilitates inference in cases with incomplete parental information, such as when one parent is specified and the other is hypothetical.

The model preserves the principles of classical Mendelian inheritance, while its probabilistic framework naturally accommodates uncertainty and certain non-Mendelian phenomena. An overview of the full range of examples is provided in Section 4, with extensions beyond Mendelian genetics discussed in detail in Section 4.2. We now proceed with a formal specification of the model and its operational structure.

### 3.1. Notation

We introduce the following notation for the genetic model:

- ℐ: Set of individuals, indexed by *i ∈ ℐ*.
- 𝒢: Set of genes, indexed by *g ∈* 𝒢, with |𝒢| = *G*.
- 𝒜_*g*_: Set of alleles for gene *g*, indexed by *a ∈* 𝒜_*g*_.
- *C* ∈ ℕ: Number of copies per gene (e.g., *C* = 2 for diploid organisms), with copy index *c* ∈ {1, …, *C*}.
- 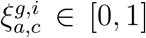: Probability that individual *i* possesses allele *a ∈* 𝒜_*g*_ in copy *c* of gene *g*, with the constraint:

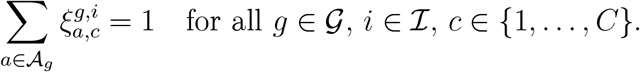
- *x*^(*i*)^: Genotypic profile of individual *i*, comprising allele distributions across all genes.
- 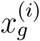: Allelic structure of gene *g* in individual *i*, given by the product over the *C* allele copies.
- 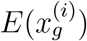: Expected allele distribution for gene *g* in individual *i*, averaged across its allele copies.
- *x*^(*i×j*)^: Genotype of an offspring resulting from the expected allele distributions of parents *i* and *j*.

### 3.2. Individual Genotype Representation

Each individual *x*^(*i*)^ is represented as a tuple of genes, with each gene defined as a product over allele distributions across its copies:

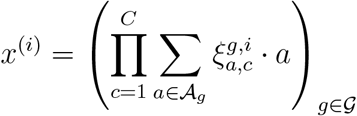

This expression represents the probabilistic presence of alleles at each gene copy, weighted by the likelihood of each allele.

### 3.3. Expected Allele Distribution

The expected allele distribution for gene *g* in individual *i*, averaged over its *C* gene copies, is given by:

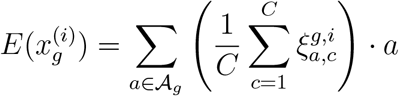

This represents the marginal distribution of alleles for gene *g*, serving as the basis for computing offspring genotypes or measuring genetic similarity.

### 3.4. Reproduction Model

To model reproduction between two individuals *x*^(*i*)^ and *x*^(*j*)^, we construct the genotype of their offspring by combining the expected allele distributions of each parent gene-wise.

For each gene *g∈* 𝒢, the offspring’s gene is represented as the product of the marginal allele distributions from both parents:

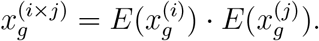

The full genotype of the offspring is then:

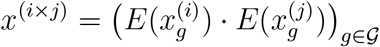

This formulation yields a joint allele distribution at each gene, representing the expected allelic composition of the offspring under the assumption of independent assortment and no recombination. The multiplication here denotes the outer product of allele distributions, producing a joint distribution over allele pairs inherited from each parent.

## 4. Examples

The following examples demonstrate how classical and non-classical genetic inheritance problems can be naturally formulated and solved using our model. While none of the examples present novel results, they serve to validate the framework by applying it to well-known scenarios from both Mendelian and non-Mendelian genetics. Our goal is to show how diverse inheritance patterns, including those involving missing data or complex genotypic interactions, can be unified under a single computational paradigm.

### 4.1. Mendelian Examples

We begin with three canonical examples from Mendelian genetics. These are chosen to illustrate increasing levels of complexity: from a basic monohybrid cross to a dihybrid test cross with missing genotype data. Through these examples, we demonstrate how the core components of our model— genotypic representation, expectation-based inheritance and inference—operate in practice.

#### 4.1.1. Monohybrid Cross

##### Example 1.

*Diploid reproduction with one gene and two alleles*

Consider a single gene *g* with alleles 𝒜_*g*_ = {*A, a*}. Individuals are diploid (*C* = 2), with each allele copy having a probability distribution over all possible alleles.

Let two individuals *x*^(*i*)^ and *x*^(*j*)^ have the following allele distributions:

- Individual *x*^(*i*)^ with genotype *Aa*:

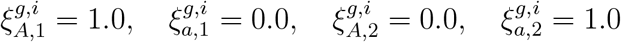

Representation:

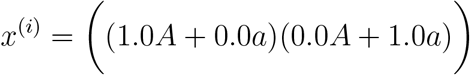
- Individual *x*^(*j*)^ with unknown genotype (equal probabilities):

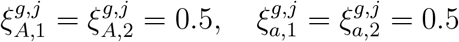

Representation:

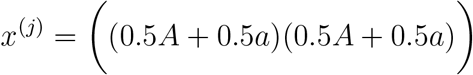

Expected allele distributions:

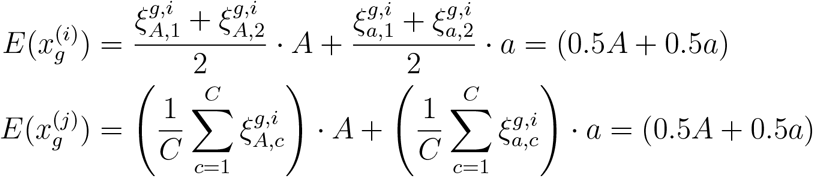

Progeny *x*^(*k*)^:

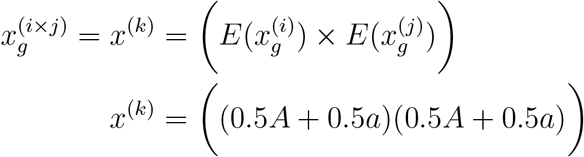

We get the following probabilities:

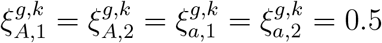

To find the probabilities of each genotype, we just need to compute the outer product of the gene.

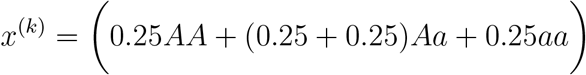

Therefore we get

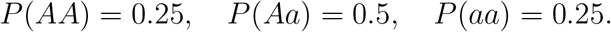

This reproduces the standard monohybrid cross outcome using our probabilistic model.

Henceforth, we will omit the explicit allele-level probability assignments (i.e. *ξ* values) and instead provide only the genotype representations of individuals in simplified form, for clarity and brevity.

#### 4.1.2. Dihybrid Cross

##### Example 2.

*Diploid reproduction with two genes and two alleles each (dihybrid cross)*

Consider genes *g*_1_ and *g*_2_ with:

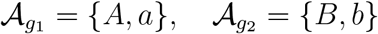

Consider two individuals,

- Individual *x*^(*i*)^ with genotype *AaBb*:

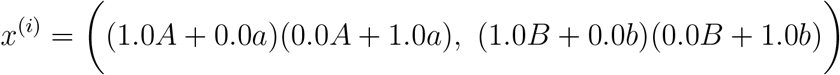
- Individual *j* with an unknown genotype at both loci:

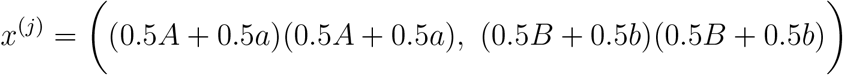

Calculating expected distributions,

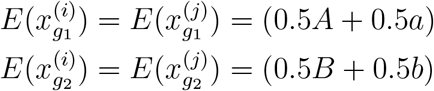

We now write the offspring distribution as

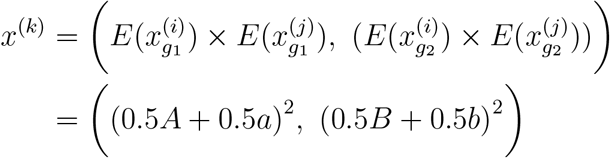

Calculating the genotypic probabilities,

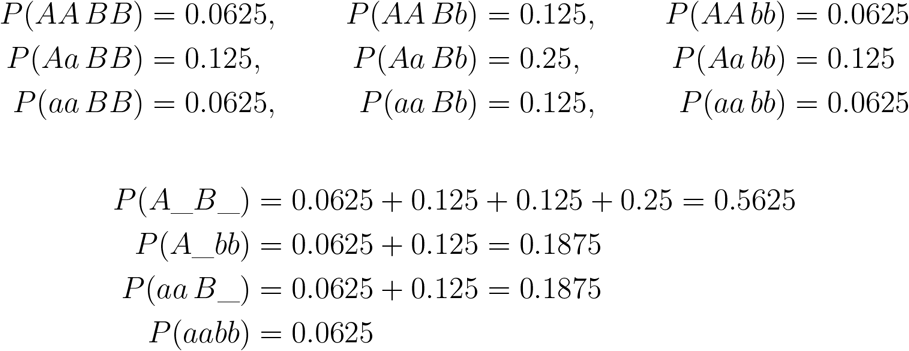

And we find the classic Mendelian dihybrid ratio of

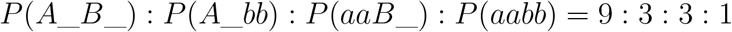

We now examine a case in which the parental genotype is partially unknown and inferred from the offspring distribution.

#### 4.1.3. Dihybrid Test Cross with Missing Alleles

##### Example 3.

*Dihybrid test cross: one unknown allele in genotype Aab*? *×aabb*

Consider genes *g*_1_ and *g*_2_ with:

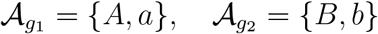

Consider two individuals:

- Individual *x*^(*i*)^ has genotype *AaB*?, where the second allele for gene *g*_2_ is unknown. Let *p* ∈ [0, 1] be the probability that the missing allele is *B*. Then the individual’s representation is:

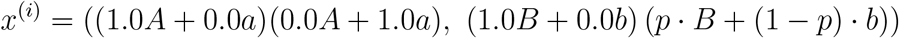
- Individual *x*^(*j*)^ has a homozygous recessive genotype at both loci:

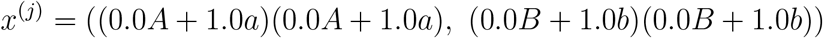

Calculating expected distributions,

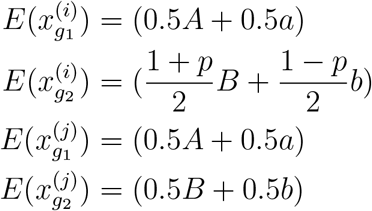

Writing the offspring distribution we get,

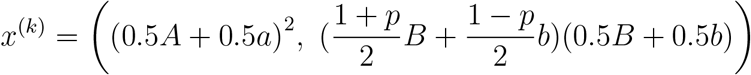

Expanding the products, we compute the genotypic probabilities for each gene independently:

For gene *g*_1_:

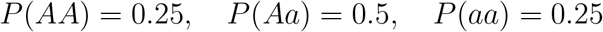

For gene *g*_2_:

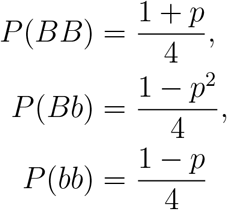

The full genotype probabilities are obtained by taking the product over gene pairs (assuming independence):

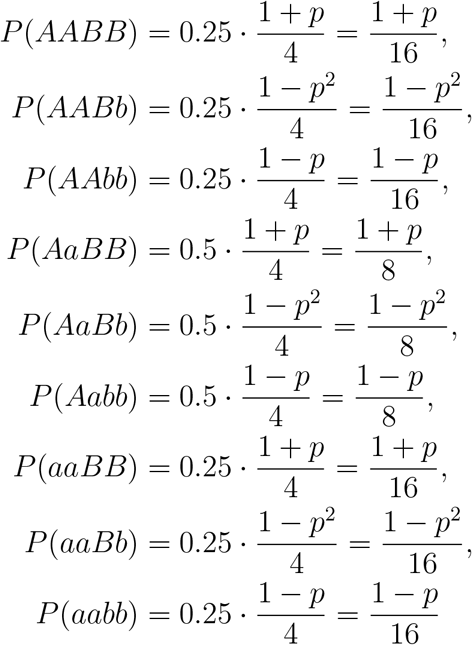

These probabilities can be compared with empirical offspring distributions to infer the likely value of *p*, completing the inference of the unknown parental allele.

This shows how the model can handle incomplete parental data while remaining consistent with classical Mendelian results.

These examples confirm that classical Mendelian problems, including those involving incomplete or missing genotypic information, are easily handled within the model. The consistency of representation across problems highlights the generality and flexibility of our approach.

### 4.2. Non-Mendelian Systems

We now turn to systems that deviate from classical Mendelian inheritance, including sex-linked traits, incomplete dominance, codominance, chromosomal abnormalities, and polygenic traits. These examples serve both to illustrate the model’s extensibility and to identify its limitations. While not exhaustive, the selected cases represent a broad spectrum of inheritance modes found in genetics.

#### 4.2.1. Sex-linked Inheritance

##### Example 4.

*Sex-linked inheritance*

One of the main challenges with sex-linked inheritance is the inherent asymmetry, which traditional genetic algebra systems are not well-equipped to handle. In our model, we address this by allowing sex to be represented probabilistically, which introduces a form of pseudo-symmetry into the system.

We first represent the sexes of individuals, without reference to any X-linked condition, as follows.

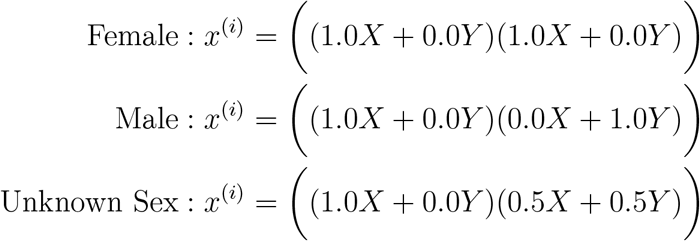

Now, consider a single gene *g* following an X-linked recessive inheritance pattern with 𝒜_*g*_ = {*X*′, *X, Y*}. Individuals are diploid and may have the following genotypes.

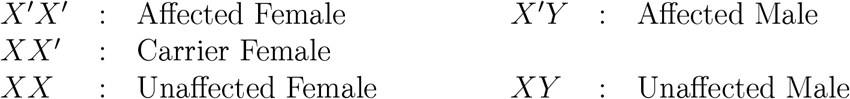

Let individual *x*^(*i*)^ be a carrier female with genotype *XX*′, and individual *x*^(*j*)^ be an unaffected male with genotype *XY*. Their allele distributions are given by:

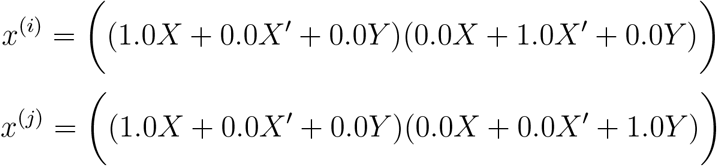

Calculating expectations and the progeny genotype,

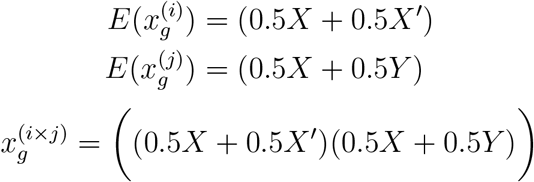

Expanding,

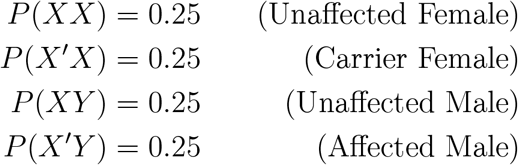

This matches classical sex-linked Mendelian ratios, now derived using allele distributions with sex represented probabilistically.

### 4.2.2. Dominance Variations

We begin with two variations on genotype-to-phenotype expression— incomplete dominance and codominance—which differ from classical dominance relationships but remain tractable under our model since the underlying genotype structure remains standard.

#### Example 5.

*Incomplete dominance in snapdragon flower colour* In incomplete dominance, heterozygous genotypes express a phenotype that is intermediate between the two homozygous conditions. Consider a gene *g* with alleles 𝒜_*g*_ = {*R, W*}, where:

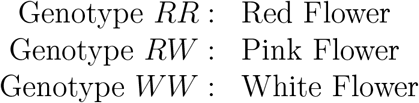

Let individual *x*^(*i*)^ be a red flower and individual *x*^(*j*)^ be a white flower:

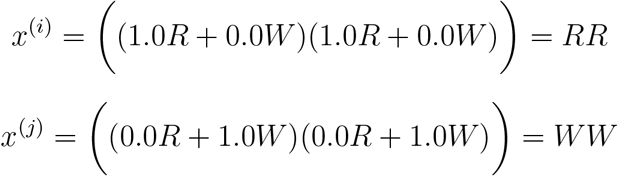

We find the progeny to have the genotype

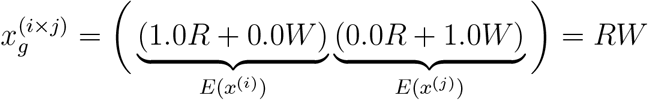

So, all the progeny will be having flowers of pink colour.

This result is fairly trivial to calculate. However, we included it to show that the model is only concerned with the genotype, and not the phenotype.

#### Example 6.

*Codominance in human blood type*

In codominance, both alleles in a heterozygous genotype are fully expressed. The ABO blood group system is a classic example, with the alleles:

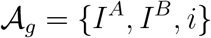

where:

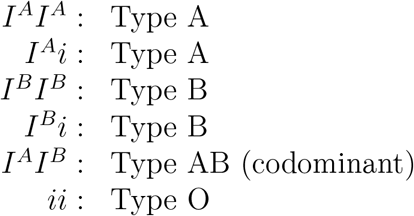

Let individual *x*^(*i*)^ have blood type A with an unknown genotype, and individual *x*^(*j*)^ have blood type B with an unknown genotype.

We know that individual *x*^(*i*)^ has at least one *I*^*A*^ while the other allele can be *I*^*A*^ or *i*. Similarly, we know the genotype of *x*^(*j*)^ contains at least one *I*^*B*^.

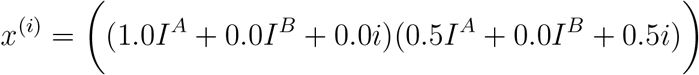

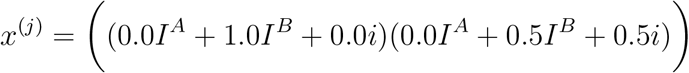

Calculating expectations,

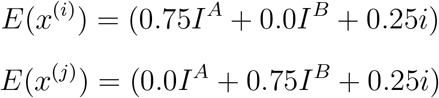

We find the progeny to have

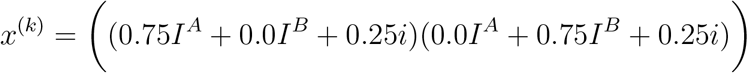

Expanding this product:

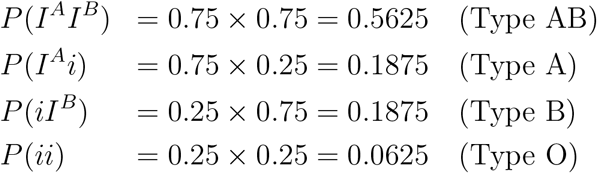

### 4.2.3. Chromosomal Abnormalities

We next examine cases involving non-standard chromosome numbers, including polyploidy and aneuploidy. These examples test the model’s ability to represent deviations in ploidy and to accommodate altered segregation patterns.

#### Example 7.

*Polyploidy: triploid inheritance*

In polyploid organisms, individuals have more than two sets of chromosomes. Let *C* = 3, representing triploidy. Consider a gene *g* with alleles 𝒜_*g*_ = {*A, a*}. Let individual *x*^(*i*)^ be a triploid gene with genotype *AAa*. We can write th allele distributions as:

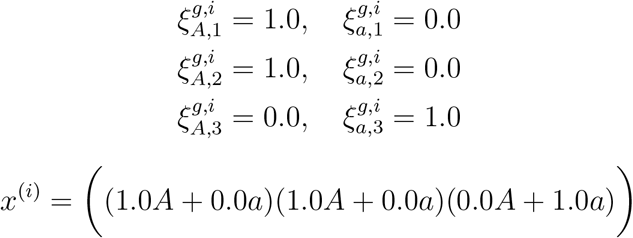

And we can calculate the expected distribution as follows:

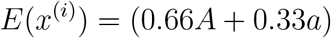

#### Example 8.

*Aneuploidy: Trisomy 21 (Down syndrome)*

Aneuploidy is the presence of an abnormal number of chromosomes. In trisomy, a chromosome appears three times instead of the normal two. Consider a simplified gene *g* on chromosome 21 with alleles 𝒜_*g*_ = {*N, n*}. Let individual *x*^(*i*)^ have three copies: *NNn*

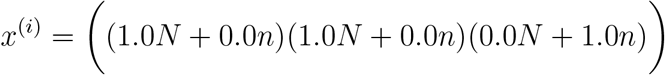

We can simply calculate expectation to be

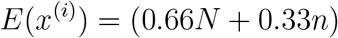

If passed to offspring, nondisjunction during meiosis may lead to gametes with:

- 2 alleles (disomic): (*E*(*x*^(*i*)^))^2^
- 1 allele (normal): *E*(*x*^(*i*)^)
- 0 alleles (nullisomic, typically nonviable): (*E*(*x*^(*i*)^))^0^

#### Example 9.

*Uniparental Disomy (UPD)*

In uniparental disomy, both copies of a chromosome (or chromosomal region) are inherited from the same parent, violating normal biparental inheritance. Let *E*(*x*^(*i*)^) be the expectation of a gene *g* in an individual 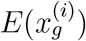. We can replicate UPD as

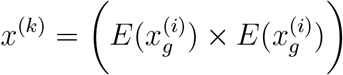

#### 4.2.4. Alternative Genetic Systems

Finally, we consider two distinct extensions: haplodiploidy, a system with fundamentally different sex determination logic, and polygenic inheritance, where multiple genes contribute to a single trait. These examples further illustrate the model’s flexibility, even in cases where the genetic architecture is more diffuse or abstract.

##### Example 10.

*Haplodiploidy in honeybee sex determination*

In honeybees, sex is determined not by sex chromosomes, but by ploidy:

- Females (workers and queens) are diploid, arising from fertilized eggs.
- Males (drones) are haploid, arising from unfertilized eggs (i.e., parthenogenesis).

Let gene *g* reside at the *csd* (complementary sex determiner) [11] locus, with allele set 𝒜_*g*_ = {*A, B, C*, …}. An individual’s sex is determined as follows:

- Heterozygous (e.g., *AB*) *→* Female
- Homozygous or hemizygous (e.g., *AA* or *A*) →Male (but diploid homozygotes are typically non-viable)

Let a queen *x*^(*q*)^ have genotype *AB* at the csd locus:

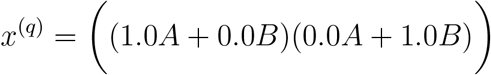

Let a drone *x*^(*d*)^ have genotype *C* (haploid):

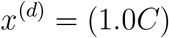

We compute expectations:

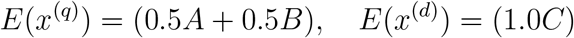

Progeny genotypes are then:

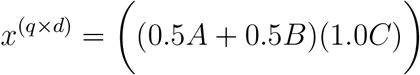

Expanding:

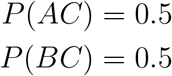

Since both *AC* and *BC* are heterozygous at the csd locus, all progeny are diploid females.

Now, for unfertilized eggs, we model parthenogenesis:

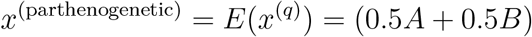

These are haploid males, with:

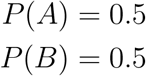

This demonstrates that the model captures the key features of haplodiploidy:

- Fertilized eggs *→* Diploid females (if heterozygous at csd)
- Unfertilized eggs *→* Haploid males

##### Example 11.

*Polygenic Inheritance: Human Skin Color*

Polygenic inheritance involves multiple genes contributing additively to a trait. Human skin color is a classic example influenced by several genes. We consider four genes known to affect pigmentation: TYR, P, SLC45A2, and MC1R [2].

Let each gene *g* ∈ {TYR, P, SLC45A2, MC1R} have an allele set 𝒜_*g*_ and an individual *x*^(*i*)^ be diploid with alleles at each locus.

For simplicity, assume:

- TYR: 𝒜_TYR_ = {*T, t*} (T: functional, t: reduced function)
- P: 𝒜_P_ = {*p*_1_, *p*_2_, *p*_3_ …} (polymorphic alleles)
- SLC45A2: 𝒜_SLC45A2_ = {*S, s*} (S: pigmented, s: less pigmented)
- MC1R: 𝒜_MC1R_ = {*M, m*} (M: eumelanin-promoting, m: pheomelanin-promoting)

Let individual *x*^(*i*)^ have the following genotypes:

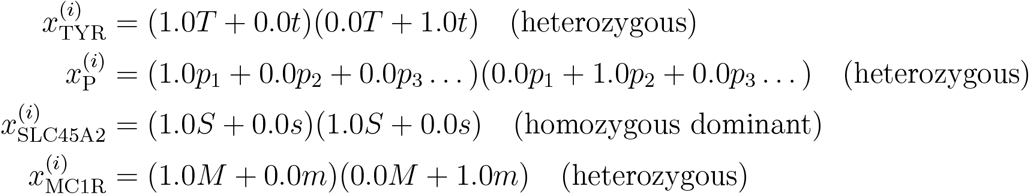

We calculate expectations for each gene:

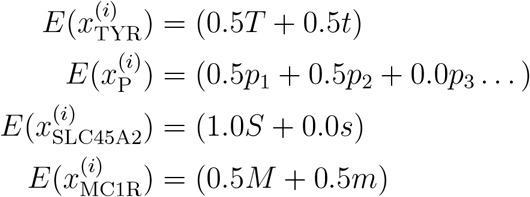

Polygenic phenotype (e.g., pigmentation level) can be modeled as an additive function of allele effects. Let *ϕ*_*g*_ : 𝒜_*g*_ → ℝ be a function assigning a pigmentation weight to each allele:

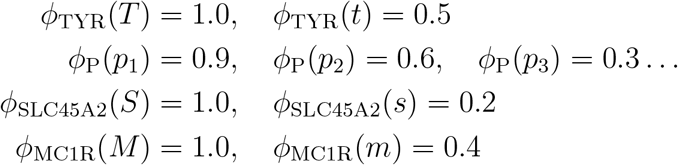

Then the expected pigmentation score is:

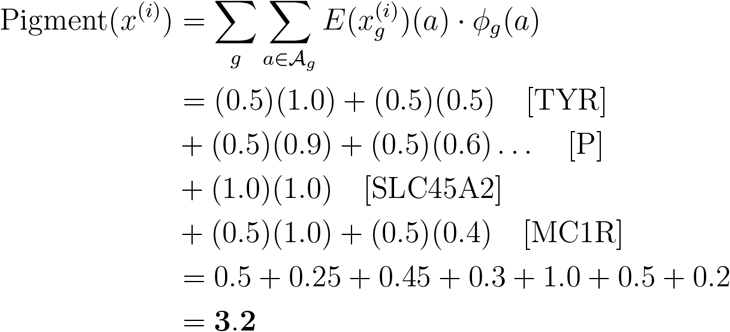

This abstract score could be normalized or mapped to skin tone categories, illustrating how multiple loci with varying alleles contribute quantitatively to pigmentation.

Together, these examples highlight the versatility of the framework in capturing a wide variety of inheritance scenarios. While certain biological processes (e.g., epigenetic modification or genomic imprinting) may fall outside the current scope, the model remains broadly applicable across classical and modern genetic problems. Its unified representation and inference capabilities offer a structured approach to reasoning about heredity in both theoretical and applied contexts.

## 5. Discussion and Limitations

The model introduced here provides a streamlined algebraic framework for simulating Mendelian inheritance across multiple genes with constant-time computational complexity. Its core innovation lies in the use of *genetic expectations*—a probabilistic analogue of expected value—that enables scalable and modular representation of genotype transmission across generations. This approach allows for efficient handling of missing or uncertain data, a frequent challenge in population-scale genetic studies.

From a practical standpoint, the model offers several key advantages. It is simple to implement, easily parallelizable, and well-suited for integration into simulation platforms, genetic screening pipelines, and educational tools. Despite its simplicity, the framework is extensible: it already accommodates basic chromosomal complexities such as polysomic inheritance and polyploidy. Validation against classical Mendelian outcomes, including monohybrid and dihybrid crosses, as well as more complex multi-gene interactions under ideal conditions, confirms its internal consistency.

However, these benefits come with certain limitations due to simplifying assumptions that prioritize computational efficiency. Chief among them is the assumption of independent assortment, which excludes genetic linkage and recombination. The model does not impose any mating structure, effectively assuming random mating in the absence of preferences or constraints, and it omits evolutionary forces such as selection, mutation, and genetic drift. Although the model is grounded in Mendelian inheritance, it is flexible enough to accommodate certain deviations. However, it does not currently support more complex phenomena such as gene conversion or epigenetic modifications. These processes involve context-dependent, often non-linear mechanisms that fall outside the model’s algebraic framework. Additionally, non-diploid inheritance is not supported.

A notable limitation is the absence of phenotype modeling. Many phenotypic traits involve complex interactions—such as epistasis (gene-gene interaction), pleiotropy (one gene affecting multiple traits), and environmental influence—that are not captured by the current framework. Incorporating these effects would require significant reformulation to support non-linear, context-dependent interactions, possibly involving external datasets or learned models.

The model also excludes positional information, which limits its ability to simulate genetic linkage. Addressing this would require incorporating chromosomal coordinates and a more sophisticated recombination mechanism, which would compromise the current model’s independence assumptions and constant-time complexity.

Other biological complexities fall outside the scope of the present framework. These include mosaicism and chimerism, where individuals harbor multiple genetic lineages, as well as de novo mutations and somatic variation. The model assumes a single, stable (though probabilistically expressed) genotype per individual, which precludes capturing such phenomena.

Finally, while illustrative examples demonstrate concordance with classical inheritance patterns, no large-scale empirical validation has yet been performed. Systematic comparison with real-world datasets or established genetic simulators is necessary to assess robustness, generalizability, and practical utility.

Future work may focus on relaxing the assumption of independence to allow modeling of linkage and recombination, incorporating limited gene-gene and gene-environment interactions, and validating the model in applied contexts such as genomic surveillance and synthetic biology. Connecting this genotype-based model to phenotype predictors or AI systems represents a promising avenue for further development.

## 6. Conclusion

We introduced a simple and efficient algebraic model for simulating Mendelian inheritance using genetic expectations. The framework supports multiple genes, handles uncertain data, and runs in constant time, making it suitable for large-scale simulations. While it simplifies some biological complexities— such as linkage, recombination, and phenotype modeling—it remains consistent with classical genetics and is easy to extend. Future work will focus on expanding the model to include more realistic genetic features and validating it with real-world data.

## Author Contributions

P. N. Bhat developed the theoretical framework, implemented the model, and wrote the initial draft of the manuscript. D. Pal supervised the project, provided critical feedback, and contributed to the manuscript’s final revision.

## Acknowledgements

P. N. Bhat gratefully acknowledges Prof. Vinay Madhusudanan, Department of Mathematics, Manipal Institute of Technology, for his insightful discussions and constructive feedback, which significantly contributed to the development of this work.

## Footnotes

2 Muller’s ratchet specifically applies to asexual reproduction, whereas Gonshor’s algebra makes no such distinction

